# Consistent Reanalysis of Genome-wide Imprinting Studies in Plants Using Generalized Linear Models Increases Concordance across Datasets

**DOI:** 10.1101/180745

**Authors:** Stefan Wyder, Michael T. Raissig, Ueli Grossniklaus

**Affiliations:** Department of Plant and Microbial Biology & Zurich-Basel Plant Science Center, University of Zurich, Zollikerstrasse 107, CH-8008 Zurich, Switzerland; Present address: Department of Biology, Stanford University, Gilbert Building, 371 Serra Mall, Stanford, CA 94305-5020.

**Keywords:** genomic imprinting, allele-specific expression, RNA-seq, consistent reanalysis, generalized linear model, Arabidopsis, maize

## Abstract

Genomic imprinting leads to different expression levels of maternally and paternally derived alleles. Over the last years, major progress has been made in identifying novel imprinted candidate genes in plants, owing to affordable next-generation sequencing technologies. However, reports on sequencing the transcriptome of hybrid F1 seed tissues strongly disagree about how many and which genes are imprinted. This raises questions about the relative impact of biological, environmental, technical, and analytic differences or biases. Here, we adopt a statistical approach, frequently used in RNA-seq data analysis, which properly models count overdispersion and considers replicate information of reciprocal crosses. We show that our statistical pipeline outperforms other methods in identifying imprinted genes in simulated and real data. Accordingly, reanalysis of genome-wide imprinting studies in *Arabidopsis* and maize shows that, at least for the *Arabidopsis* dataset, an increased agreement across datasets can be observed. For maize, however, consistent reanalysis did not yield in a larger overlap between the datasets. This suggests that the discrepancy across publications might be partially due to different analysis pipelines but that technical, biological, and environmental factors underlie much of the discrepancy between datasets. Finally, we show that the set of genes that can be characterized regarding allelic bias by all studies with minimal confidence is small (~8,000/27,416 genes for *Arabidopsis* and ~12,000/39,469 for maize). In conclusion, we propose to use biologically replicated reciprocal crosses, high sequence coverage, and a generalized linear model approach to identify differentially expressed alleles in developing seeds.

## INTRODUCTION

In a diploid cell, the maternal and paternal alleles of a given gene usually share the same expression state in a specific tissue, meaning that they are either both expressed or both silent. Important exceptions to this rule are genes regulated by genomic imprinting, where the expression state depends on the parental origin of the alleles and only one is expressed while the other remains silent or is weakly expressed. The two alleles do not differ in their sequence but rather carry parent-specific, epigenetic imprints that allow the cell to distinguish the two alleles (Reik and Walter, 2001; Grossniklaus, 2007; Bartolomei and Ferguson-Smith, 2011; Ferguson-Smith, 2011; Barlow, 2011; Raissig et al., 2011; Jiang and Köhler, 2012; Gehring, 2013). Genomic imprinting evolved independently in mammals and in flowering plants (angiosperms) (reviewed in Messing and Grossniklaus, 1999; Baroux et al., 2002; Feil and Berger, 2007; Köhler and Weinhofer-Molisch, 2010; Raissig et al., 2011; Gutierrez-Marcos et al., 2012; Pires and Grossniklaus, 2014; Rodrigues and Zilberman, 2015). In both groups, the offspring develop within the mother and depend solely on her nutrients for growth and development. This common reproductive strategy results in an intragenomic parental conflict over resource allocation, which likely underlies the evolution of genomic imprinting, at least for loci that control growth (Haig and Westoby, 1989; Moore and Haig, 1991; Pires and Grossniklaus, 2014). Accordingly, some imprinted genes in both, mammals and plants, have a role in controlling growth (e.g. Lau et al., 1994; Ludwig et al., 1996; Grossniklaus et al., 1998; Kinoshita et al., 1999; Kiyosue et al., 1999; Luo et al., 2000; Ingouff et al., 2005; Tycko and Morrison, 2002; Monk, 2015). Consistent with this function, many imprinted genes are preferentially expressed in the tissues that support embryonic growth, i.e. the placenta in mammals or the triploid endosperm in the seeds of flowering plants.

In the last decade, the advent of Next-Generation Sequencing (NGS) allowed (nearly) genome-wide imprinting studies by sequencing the transcriptome of hybrid F1 seed tissues: Given exonic polymorphisms between the parents, reads overlapping heterozygous SNPs can be assigned to their parent-of-origin, and reciprocal crosses allowed the discrimination between parent-of-origin-dependent and strain-specific genetic effects. Accordingly, a number of research groups performed genome-wide, allele-specific transcriptome profiling studies of hybrid seeds in *Arabidopsis* and maize to identify genes that are preferentially expressed from one parental allele (Autran et al., 2011; Gehring et al., 2011; Hsieh et al., 2011; McKeown et al., 2011; Waters et al., 2011; Wolff et al., 2011; Zhang et al., 2011; Pignatta and Gehring, 2012; Waters et al., 2013; Xin et al., 2013; Zhang et al., 2014; Pignatta et al., 2014). As a result, the total number of imprinted genes increased from around 20 (Raissig et al., 2011) to over 900 potentially imprinted plant genes (Gehring et al., 2011; Hsieh et al., 2011; McKeown et al., 2011; Waters et al., 2011; Wolff et al., 2011; Zhang et al., 2011; Waters et al., 2013; Xin et al., 2013; Pignatta et al., 2014).

However, comparisons of the identified imprinted candidate genes revealed little overlap between the studies (McKeown et al., 2011; Köhler et al., 2012; Pignatta and Gehring, 2012). In general, the analysis of RNA-sequencing (RNA-seq) data to identify allele-specific expression is prone to false positives due to both, biological and technical variation (DeVeale et al., 2012; Wang and Clark, 2014; Castel et al., 2015). Thus, even studies with seemingly similar design heavily disagree on the number of imprinted genes in the mouse brain, e.g. ranging from less than 200 (DeVeale et al., 2012) to over a thousand (Gregg et al., 2010). To date, although guidelines for the analysis of allele-specific expression have recently become available (Castel et al., 2015), many different methods have been applied to filter, normalize, and statistically assess allelic imbalance from RNA-seq data. For the analysis of allele-specific expression several analysis methods and software (Castel et al., 2015) have been developed, yet only very few are suitable for an analysis of imprinted expression. Moreover, no specialized method is available for statistical testing of imprinting in the triploid endosperm, where the expected allelic ratio is 2:1 because the mother contributes two genomes to this tissue. In plants, many authors have used count tests (such as Chi-Square, binomial, or Fisher’s exact tests), which heavily underestimate the count dispersion typically observed in RNA-seq data (Zhou et al., 2011; Wang and Clark, 2014; Castel et al., 2015), resulting in increased numbers of false positives particularly for large counts. Highly expressed transcripts may appear imprinted with high statistical significance, as count tests are sensitive to very small allelic imbalance at high counts, requiring additional filtering with somewhat arbitrary imbalance cut-offs.

Here, we present a new statistical approach to call imprinted genes from large allele-specific RNA-seq datasets from endosperm that outperforms other methods in simulated and real data. We propose a generally applicable approach using generalized linear models (GLM) implemented in edgeR (Robinson et al., 2010), which is based on the negative binomial distribution to deal with potential count overdispersion (Anders and Huber, 2010) as it is typically seen in RNA-seq data. The presented pipeline outperforms other methods using simulated data. Furthermore, we reanalyze the raw data from seven studies to assess the relative importance of differences in data generation and data analysis. The consistent reanalysis by the proposed pipeline results in a larger overlap of imprinted candidate genes across *Arabidopsis* datasets, but showed little improvement across maize datasets. In conclusion, consistent data analysis can improve concordance between datasets but biological and technical variation in data generation contributes most to differences in datasets.

## RESULTS

### Comparison of Genome-wide Imprinting Studies in Plants: Biological, Technical, and Statistical Differences

To shed light onto discrepancies between published studies and potential biases, we compared the genome-wide, allele-specific transcriptome profiling studies of hybrid seeds in *Arabidopsis* and maize that were designed to identify imprinted candidate genes in the endosperm. All studies were based on reciprocal crosses of two polymorphic inbred strains, and all studies used manually dissected endosperm. In *Arabidopsis*, a total of 343 genes were proposed to be maternally expressed imprinted genes (MEGs) in at least one study using the Landsberg *erecta* (L*er*) and Columbia-0 (Col-0) accessions (Fig. 1). The three studies proposed between 116 and 165 MEGs (Gehring et al., 2011; Hsieh et al., 2011; Pignatta et al., 2014). The large majority of genes (83%) were unique to a single study and only eleven MEGs were identified as imprinted in all three studies (3%; Fig. 1). 68 genes were proposed to be paternally expressed imprinted genes (PEGs) in at least one study with six PEGs being commonly identified in all three studies (9%; Fig. 1). In maize, the four available studies using the inbreds B73 and Mo17 (Waters et al., 2011; Zhang et al., 2011; Xin et al., 2013; Zhang et al., 2014) listed 322 different MEGs and 228 PEGs (Fig. 1). The majority of genes (79% and 52% for MEGs and PEGs, respectively) was proposed by a single study only. The overlap between studies was also small, with 14 MEGs (4%) and 23 PEGs (10%) being commonly identified by all four studies.

**Figure 1.**
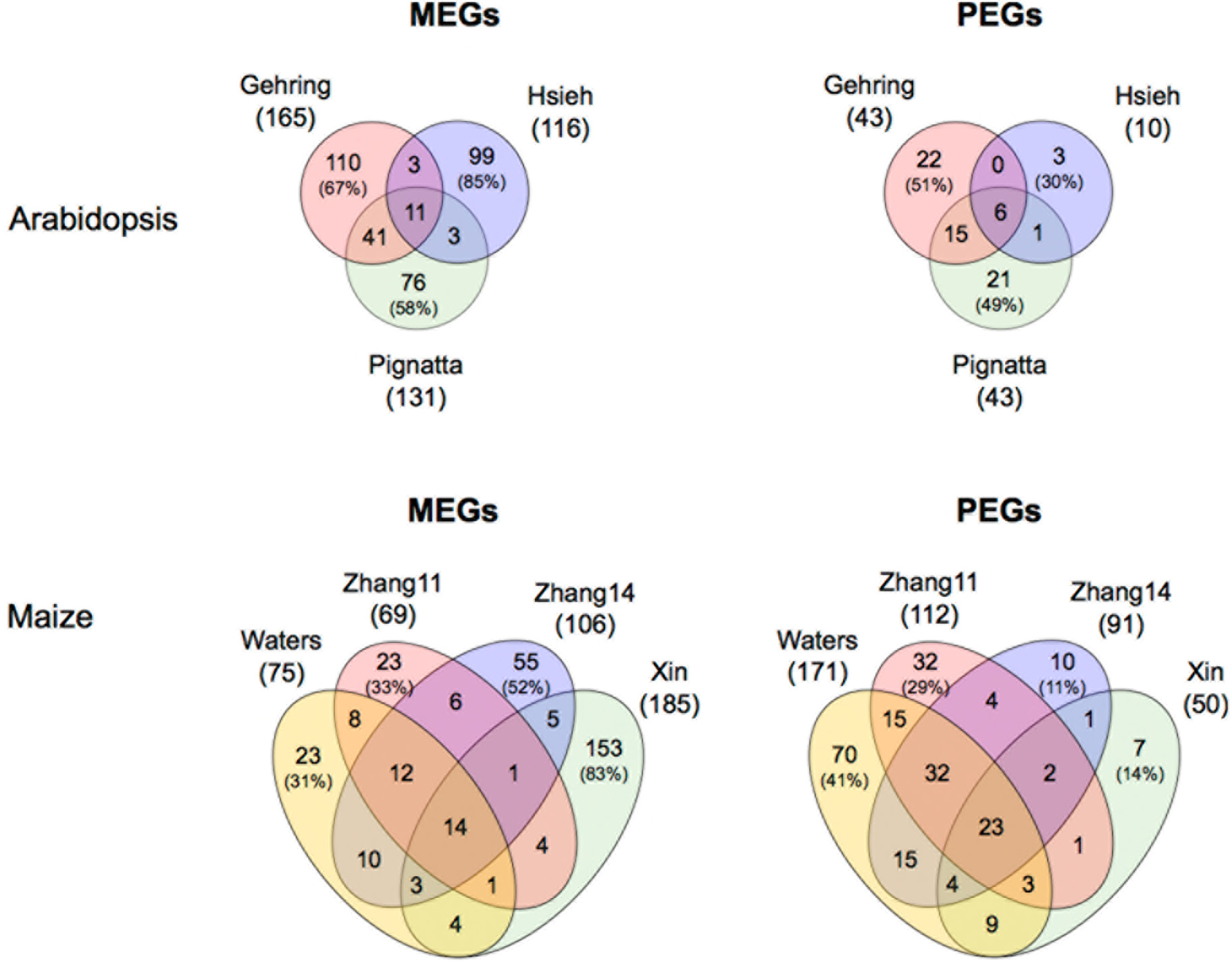
Venn diagrams showing the number of imprinted genes in hybrid endosperm reported by different studies in *Arabidopsis* L*er*/Col accessions and in maize B73/Mo17 inbreds. Numbers in brackets denote the percentage of non-shared genes relative to the full set reported. The “Xin” and "Zhang11" sets comprise genes identified at 10 days after pollination and "Zhang14" comprises genes at 12 days after pollination. Accession/inbred-specific imprinted genes were excluded from the analysis.

The low concordance between studies could be due (i) to intrinsic, biological differences (e.g. developmental stage analyzed), (ii) to technical differences (particularly library preparation and complexity, sequencing depth and batch effects, reviewed in [Wang and Clark, 2014; Castel et al., 2015]), and/or (iii) to the varying bioinformatics/statistical analysis protocols applied. As summarized in Table 1, all studies differ in terms of developmental stage and some use different accessions creating considerable biological variation. Technical differences like library preparation, sequencing platform, read length, and single-end vs. paired-end sequencing introduce a further level of variation. Particularly, the observed differences in sequencing depth, expected read-mapping biases (Degner et al., 2009), as well as in the completeness and quality of available SNP annotations, present likely technical sources of inconsistency (Table 1). Regarding statistical analysis, all studies applied count statistics (Table 1), which do not properly model count dispersion of RNA-Seq data, resulting in increased numbers of false positives particularly for large counts (Zhou et al., 2011; Wang and Clark, 2014). Lastly, the studies apply very different criteria for filtering potentially imprinted genes according to the allelic bias (Table 1). The requirement to call a gene’s expression parentally biased differed tremendously between the studies and ranged from 90% of all reads that have to derive from one parent (Waters et al., 2011), over 5 times more reads from one parent (Zhang et al., 2011), to simply assessing deviations from the expected 2:1 ratio in the endosperm (Gehring et al., 2011).

**Table 1.**
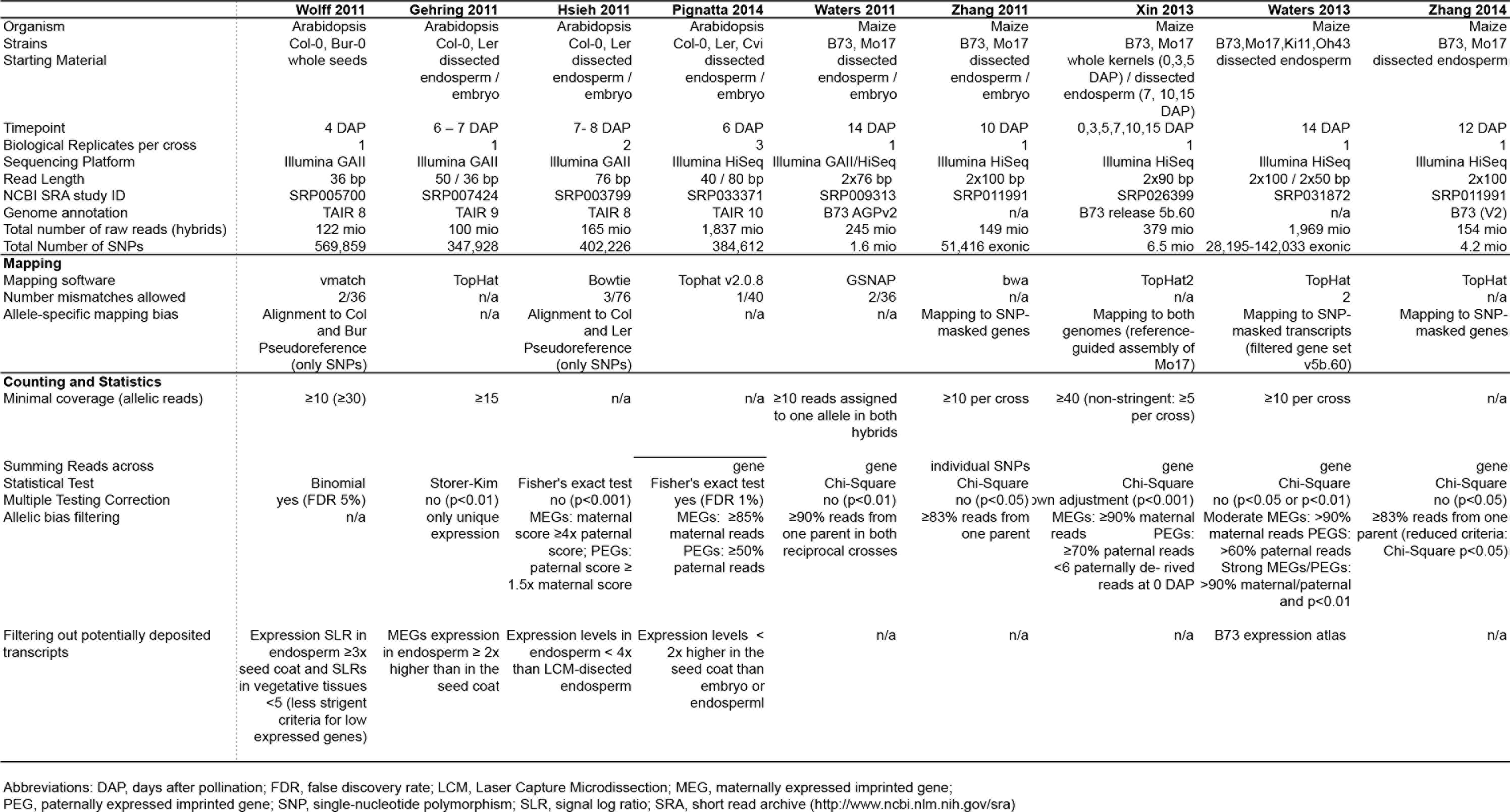
Characteristics of data generation and data analysis of published genome-wide imprinting datasets in *Arabidopsis* and maize.

### Analysis of Allelic Bias Using edgeR Outperforms other Methods

It was previously noticed that a large part of the differences between publications is owed to different statistical pipelines to call imprinted genes. When the two *Arabidopsis* datasets that analyzed the same accessions and a similar developmental stage and tissue (Gehring and Hsieh datasets) were analyzed in the same way, the overlap increased substantially (from 14 to 56 MEGs and from 6 to 18 PEGs; Gehring et al., 2011). Therefore, we created a statistical pipeline to identify genes with statistically significant allelic imbalance from the expected, endosperm-specific 2:1 ratio. Our pipeline is based on edgeR (Robinson et al., 2010) and analyzes counts by a generalized linear model (GLM), based on a negative binomial distribution in a paired design (parentals of the same cross) with two or more biological replicates (or reciprocal crosses). Importantly, edgeR models count overdispersion as shown exemplarily for the Pignatta dataset (Supplemental Figure S1).

We then tested the performance of our statistical pipeline compared to other methods based on synthetic data, where we could control the settings and the true genomic imprinting status of each gene. We simulated counts using negative binomial distributions, with mean and dispersion parameters estimated from real data (see Material and Methods). Imprinted genes were introduced by adding 200 genes with a parent-of-origin specific allelic bias: 50 MEGs each with strong (99% maternal reads) or moderate (85% maternal) allelic bias as well as 50 PEGs each with strong (34% maternal reads) or moderate (48% maternal reads) allelic bias. Random allelic read sampling was modeled by sampling from a binomial distribution.

We evaluated the performance of three different methods: our edgeR-based pipeline, Fisher’s exact test, and Stouffer’s method. Fisher’s exact test is the analysis method used in most published plant studies where each reciprocal cross is tested individually, resulting in two p-values per gene. With this method, a gene is called imprinted if both adjusted p-values independently reach significance. From now on this method will be called “Fisher-separate”. Stouffer’s method (Stouffer et al., 1949) is a method to combine p-values bearing the same null hypothesis. We combined the two p-values per gene calculated by Fisher’s exact tests using Stouffer’s method and calculated a combined adjusted p-value. From now on this method will be called "Fisher-combined".

In our simulation setting, our pipeline (edgeR) identified the largest number of observed true positives (spiked-in MEGs and PEGs) (Fig. 2A), identifying 131 true positives (of 200, 66% true positive rate [TPR]) with 8 false positives (falsely detected as imprinted), whereas the second best method, Fisher-combined, identified only 93 true positives (47% TPR) with 0 false positives (0% FPR). Fisher-separate identifies only 33 true positives (17% TPR) with 0 false positives (0% FPR). The Venn diagram (Fig. 2A) shows that 33 genes are identified between all three methods, 60 are shared between edgeR and Fisher-combined, 46 genes are only identified by edgeR and 69 true positive genes were not identified by any method. Considering the genes that were simulated to be imprinted as the true positive group and the remaining genes as the true negative group, we computed the false positive rate and the true positive rate for all possible score thresholds and constructed a ROC (Receiver Operating Characteristic) curve for each method (Fig. 2B). edgeR and Fisher-combined had a performance advantage in detecting the spike-in genes over Fisher-separate, which underperforms over the entire range. We divided the assessed genes into four equally sized bins according to their number of counts. As expected, the true positive rate (TPR) increased with larger counts per gene for all methods (Fig 2C). TPR was highest for edgeR in all categories with a large advantage in the two categories with the lowest counts, while Fisher-separate method only passed the 50% TPR for the quarter of genes with the largest counts. True positive rates also depended on the degree of allelic imbalance: at a 5% False Discovery Rate (FDR) cut-off, edgeR achieved TPRs of 80% and 54%, respectively, for strong and weak MEGs, and TPRs of 70% and 58%, respectively, for strong and weak PEGs. In contrast, the second best method Fisher-combined achieved TPRs of 60% and 42%, respectively, for strong and weak MEGs, and TPRs of 58% and 26%, respectively, for strong and weak PEGs. Simulations with count distributions similar to the largest observed datasets showed that edgeR still performed better than the other methods (data not shown).

**Figure 2.**
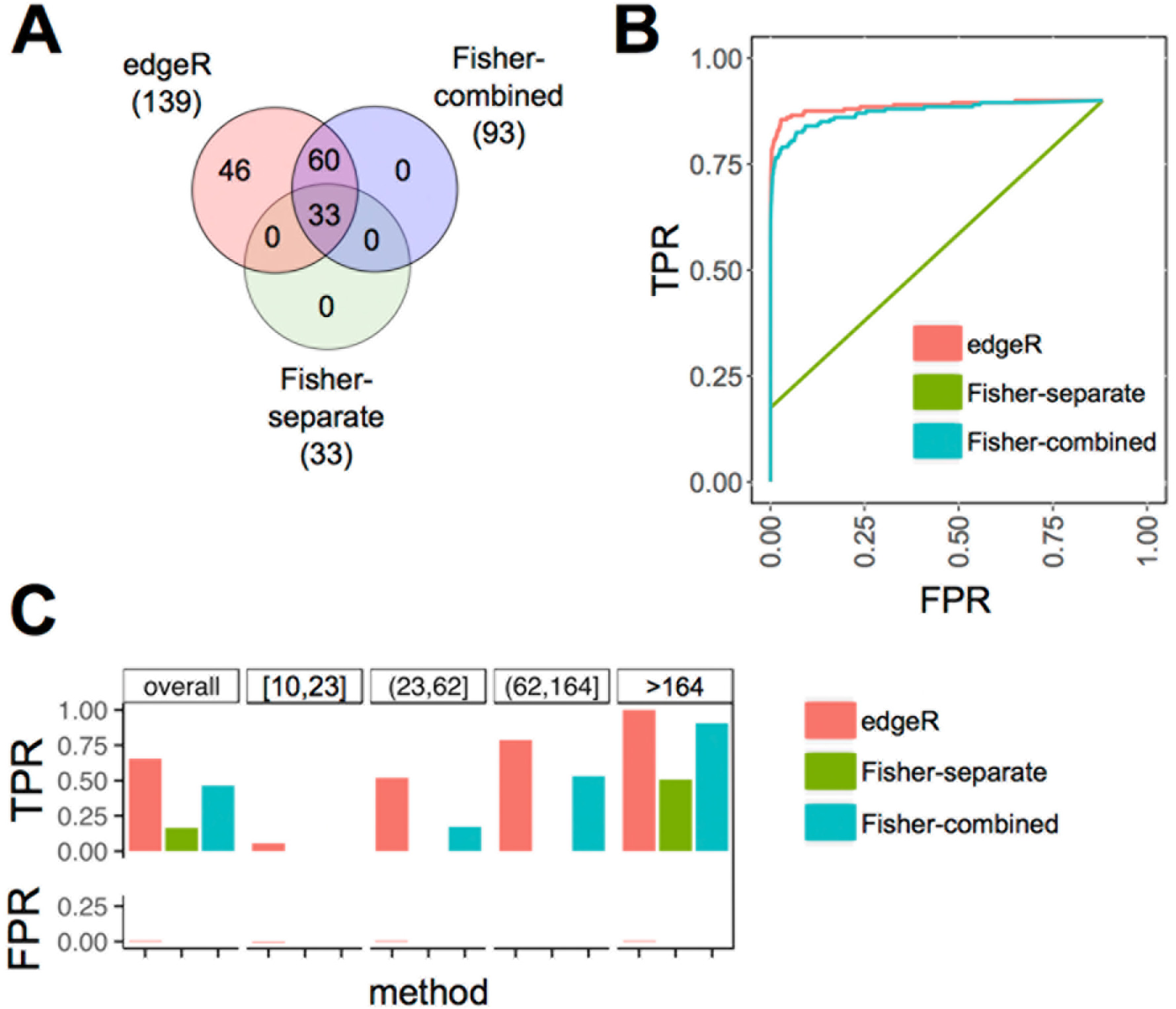
Benchmarking of three tested methods to identify imprinted genes using simulated data. A, Overlap of detected spike-in imprinted genes between different methods. B, ROC curves. C, True positive rates (TPR) and false positive rates (FPR) across four equally sized categories of genes with increasing number of counts.

### Consistent Reanalysis of Published Imprinting Studies with edgeR Identifies More Common Imprinted Candidate Genes in *Arabidopsis* but not in Maize

To test our statistical pipeline on real data, we reanalyzed the raw data from the nine published studies starting from the raw reads. We examined all reads overlapping with previously known exonic SNPs (see Materials and Methods), separated and counted maternal and paternal reads overlapping SNPs, and summed up informative (i.e. SNP containing) reads across transcripts. After discarding very lowly expressed genes (< 10 counts per gene), median reads per transcript were 95-654 for the different datasets. We then identified genes with statistically significant allelic imbalance from the expected, endosperm-specific 2:1 ratio using edgeR (Robinson et al., 2010). For simplicity, we only compared studies using L*er* and Col-0 accessions (*Arabidopsis*) or B73 and Mo17 inbreds (maize). We identified between 28-771 candidate MEGs and 33-311 candidate PEGs for the examined datasets using a 5% FDR cut-off (Fig. 3). Diagnostic plots exemplary for the Pignatta dataset are shown in Supplemental Figure S2 and display a good model fit. A list of the called imprinted candidate genes in *Arabidopsis* and maize can be found in Supplemental Table S1 and S2, respectively.

**Figure 3.**
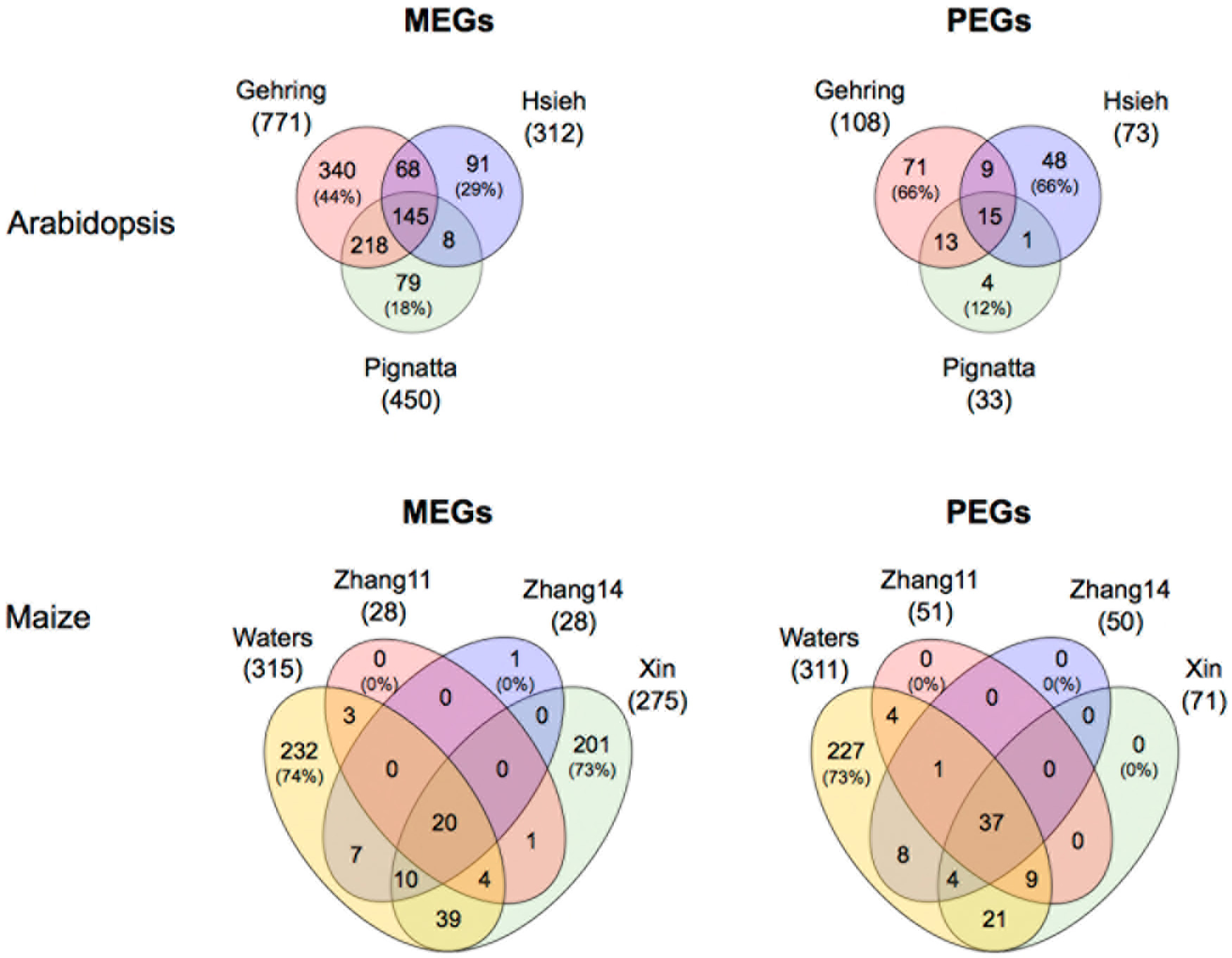
Venn diagrams showing the overlap between imprinted candidate genes across datasets when reanalyzing the raw data using the same standardized method using generalized linear models/edgeR at a FDR cut-of 5%. Numbers in brackets denote the percentage of non-shared genes relative to the full set detected in the dataset. The “Xin” and "Zhang11" sets comprise genes identified at 10 days after pollination and "Zhang14" comprises genes at 12 days after pollination.

In *Arabidopsis*, we identified a larger number of potentially imprinted genes with clearly increased overlaps: 145 common MEGs were found from all three datasets (46% of the smallest dataset [Hsieh], and 19% of the largest dataset [Gehring]) and, in addition, 68 MEGs (22% of the smaller dataset) that were shared by the Gehring and Hsieh datasets, and 218 MEGs (48%) shared by the Gehring and Pignatta datasets, which used a similar developmental stage and were generated by the same authors (Fig. 3). Now only 44% and 29% of identified MEGs were unique to the Gehring and Hsieh datasets, respectively. In the Pignatta dataset, only a small minority of MEGs (79/450, 18%) and PEGs (4/33, 12%) were not shared with any other dataset, potentially owing to the fact that (i) the Pignatta dataset had the largest sequencing depth and therefore the largest gene coverage, and (ii) three biological replicates per cross could be analyzed, allowing the identification of high-confidence candidates with a small number of false positives. We also identified an almost three-fold increased number of PEGs (186 instead of 68 genes as originally published), although with an increased proportion of non-shared PEGs in the Gehring (from 51% to 66%) and Hsieh (from 30% to 66%) datasets.

In maize, we identified a large number of imprinted candidate genes, 518 MEGs and 311 PEGs (Fig. 3). 85 MEGs (16%) and 84 PEGs (27%) were identified from at least two different datasets, leading to an overlap of MEGs similar to the one found in the originally published lists (Fig. 3). The number of candidate imprinted genes varied a lot between datasets, limiting the maximum number of genes shared by all four datasets. The proportion of non-shared MEGs and PEGs decreased for all datasets, except for Waters MEGs and PEGs which increased from 31% to 74% and from 41% to 73%, respectively, likely due to the significant increase in the number of imprinted candidate genes (Fig. 3). In contrast, the Zhang datasets, as well as the PEGs identified from the Xin dataset, showed almost no exclusive candidate imprinted genes.

Furthermore, we identified the top50 imprinted candidate genes, which are statistically most significant for each dataset and compared the number of shared genes (Supplemental Figure S3). Between 13-28 candidate genes were common to all compared datasets and the proportion of non-shared candidates decreased or was similar as in the originally published lists except for the Hsieh PEGs.

In summary, a reanalysis of the datasets using our edgeR-based pipeline produced gene lists with a clearly larger overlap in *Arabidopsis*, despite identifying a larger number of imprinted candidate genes with less pronounced allelic imbalance. Thus, the Jaccard similarity indices between *Arabidopsis* datasets were higher both for MEGs and PEGs after reanalysis with edgeR compared to the originally published gene lists (Supplemental Table S3). For maize, however, we did not see an increase in dataset concordance after reanalysis.

### Reanalysis Using the edgeR Analysis Pipeline Identifies Novel Imprinted Candidate Genes in Comparison to the Original Analysis

Having identified imprinted candidate genes using our new analysis pipeline, we compared the candidate genes pairwise with the lists from the original publications (Fig. 4). When comparing with our candidate MEGs and PEGs at a 5% FDR cut-off, 26-95% of previously published imprinted candidate genes were also identified by our pipeline.

**Figure 4.**
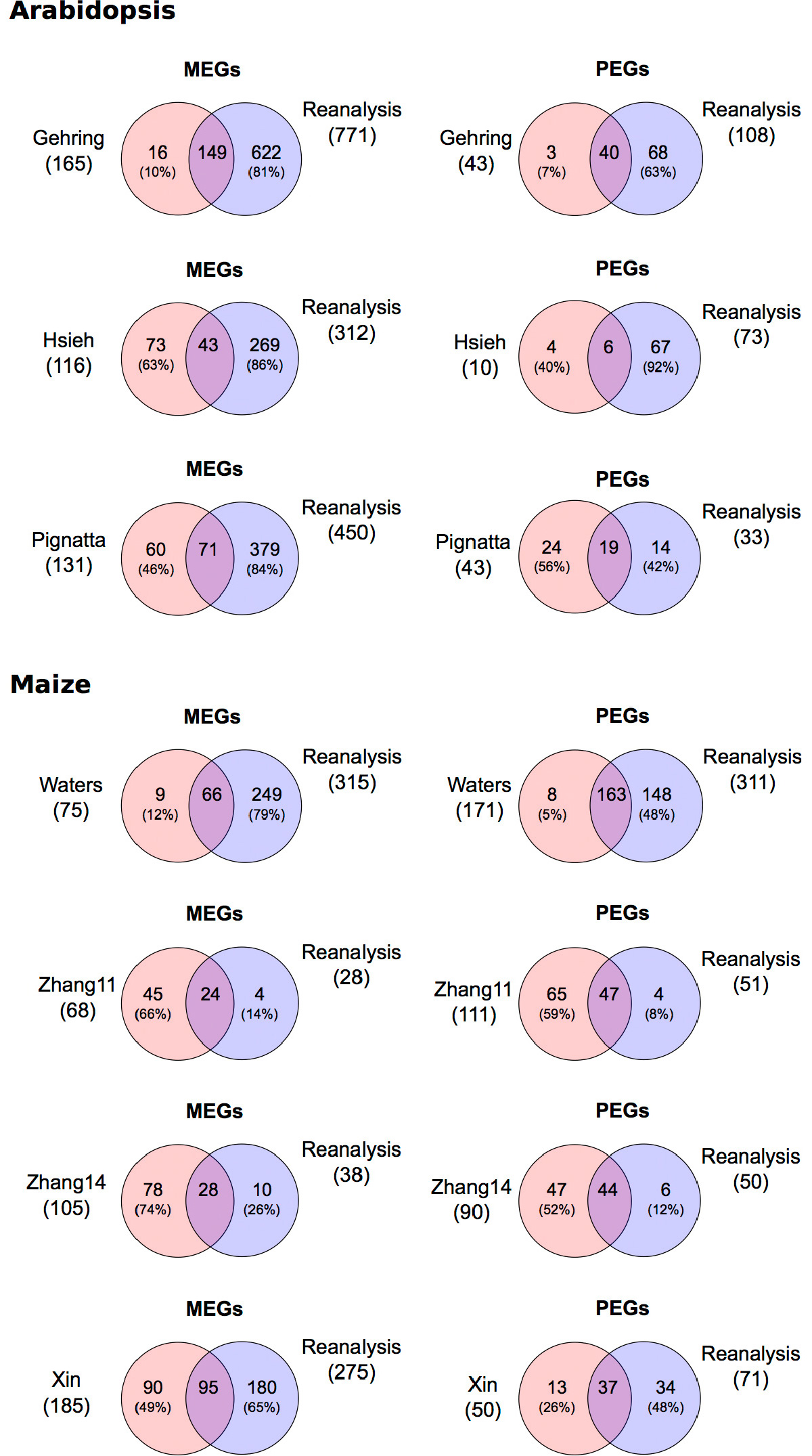
Pairwise comparison of imprinted genes between the originally published analysis and the reanalysis using generalized linear models and edgeR with a 5% FDR cut-off. Numbers in brackets denote the percentage of non-shared genes relative to the full set detected in the dataset. For the “Xin” dataset only one timepoint (10 days after pollination) is shown. The "Zhang11" set comprises genes identified at 10 days after pollination and "Zhang14" comprises genes expressed at 12 days after pollination.

From the reanalysis we selected the genes with topmost significance to get the equivalent number of the previously published imprinted candidate genes and compared them to the original publications. *Arabidopsis* PEGs and maize MEGs and PEGs generally overlapped at 40-78% between the reanalysis and the original gene lists (Supplemental Figure S4), much higher than for *Arabidopsis* MEGs, where often only negligible overlaps were observed.

In conclusion, the reanalysis with a standardized analytical pipeline identifies a large number of similar candidates but also additional novel candidate genes. Furthermore, it fails to identify previously called imprinted candidates, likely owing to the improved statistical analysis, which takes into account count overdispersion and neglects large allelic biases in transcripts covered by a low number of reads.

### Consistent Reanalysis Using Fisher’s Exact Test Identifies Fewer Overlapping Imprinted Candidate Genes

We also reanalyzed the data from the seven studies using the "Fisher-combined" method in order to assess its performance with real data. We selected the genes with topmost significance to get for each dataset the equivalent number of the previously published imprinted candidate genes (Supplemental Figure S5). In *Arabidopsis*, the number of genes identified from all three datasets (5 MEGs and 0 PEGs; Supplemental Figure S5) was only a fraction than after reanalysis with edgeR (58 MEGs and 8 PEGs; Supplemental Figure S6).

Also the number of genes shared between at least two different datasets (60 MEGs and 6 PEGs; Supplemental Figure S5) was clearly smaller than after reanalysis with edgeR (123 MEGs and 27 PEGs; Supplemental Figure S6). The proportion of non-shared genes was 50-95% per dataset, much higher than after reanalysis with edgeR, where proportions of non-shared genes were 10-40%. A reanalysis of the maize datasets using the "Fisher-combined" method produced a similar concordance between datasets as edgeR (Supplemental Figures S5 and S6). Accordingly, Jaccard similarity indices between datasets were markedly higher in *Arabidopsis* after reanalysis with edgeR both for MEGs and PEGs compared with Fisher-combined reanalysis or the originally published gene lists (Supplemental Table S3).

### Power of Detecting Imprinted Genes Is Relatively Small due to Non-saturating Sequencing Depth

The power of allelic imbalance detection depends mainly on the degree of allelic bias, the sequencing read length and coverage (expression strength), as well as the divergence of the crossed strains (number of SNPs per transcript) (Fontanillas et al., 2010). Although the three *Arabidopsis* studies all use the same accessions, the number of callable genes ranges from 8,319 for the Hsieh dataset to 14,229 for the Pignatta dataset (Fig. 5A): Among the 27,416 protein-coding *Arabidopsis* genes 31% do not overlap with any exonic SNP and their imprinting cannot be assessed. Another 17-39% of the genes are not or only very weakly expressed (<10 allelic counts overall), and were thus discarded from further analysis. Ultimately, only 30-52% of all predicted *Arabidopsis* genes can be assessed for genomic imprinting in the three different studies as only those have a sufficient number of allele-specific reads to identify statistically significant biased expression (Fig. 5A). 7,876 genes (29% of all *Arabidopsis* genes) can be assessed for genomic imprinting by all three datasets (Fig. 5B).

**Figure 5.**
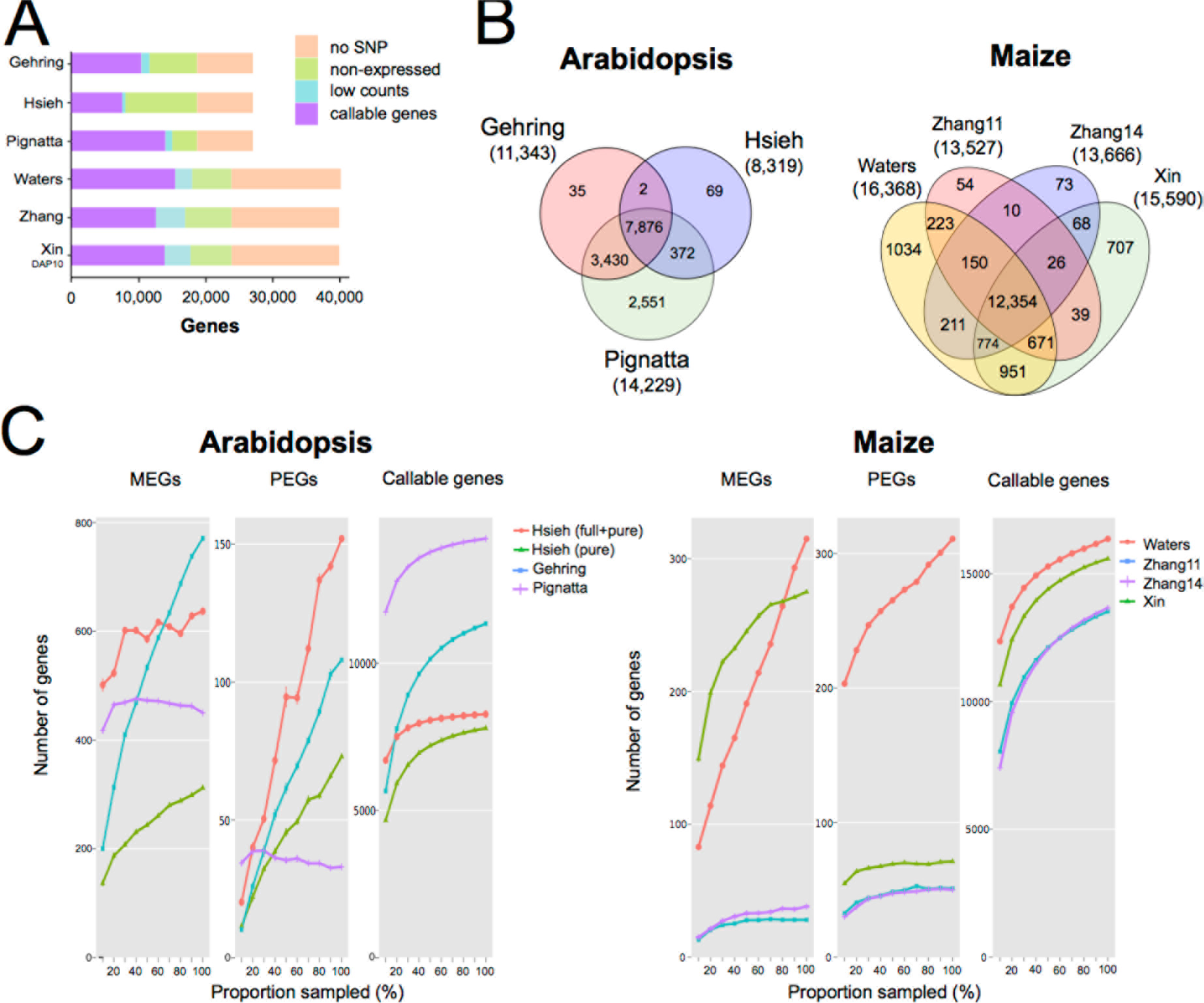
Power of detecting imprinted candidate genes in reanalyzed datasets. A, Callable genes assessable for genomic imprinting were required to overlap with at least 1 exonic SNP and to have read counts of at least 10. Number of genes are shown as percentages of the total number of genes. B, Venn diagrams showing the overlap of callable genes for *Arabidopsis* and maize datasets. C, Saturation curves showing numbers of detected MEGs and PEGs from various datasets. The curves were generated by randomly sampling increasing proportions of each dataset and identifying imprinted candidate genes using the same pipeline. Values are means (and standard errors) of 10 random subsamples.

Among the 39,469 maize protein-coding genes, 33% do not overlap with any exonic SNP (Fig. 5A) between the B73 and Mo17 inbreds. Another 26-33% were not or very weakly expressed (<10 counts overall) and were discarded (Fig. 5A). In the end, between 34% (Zhang11) and 41% (Waters) of maize genes can be assessed for genomic imprinting per dataset and only 31% of all maize genes can be assessed in all datasets (Fig. 5A and 5B). In conclusion, even after consistent reanalysis, the set of genes that can be assessed for genomic imprinting differs considerably between the datasets.

In addition, sequencing depth is far from being saturated, as shown by random subsampling of each dataset and detecting MEGs and PEGs by our edgeR-based pipeline (Fig. 5C). For most datasets, the number of detected MEGs and PEGs do hardly flatten with increasing sampling proportions. The relatively flat slopes for the total number of callable genes indicate that a huge increase in sequencing depth would be required to assess the remaining 4-10% genes that were expressed, but did not reach the minimal read coverage of 10 reads.

## DISCUSSION

### Reanalysis of Imprinting Studies Reveals that Biological and Technical Differences Strongly Contribute to Biases in Identifying Imprinted Genes

Published studies strongly disagree in the extent and composition of imprinted genes in the endosperm (Fig. 1). We reanalyzed both simulated and real data from seven genome-wide imprinting studies in *Arabidopsis* and maize, using a standardized bioinformatics pipeline based on generalized linear models. In *Arabidopsis*, our analysis identified an increased number of imprinted genes co-identified by at least two different datasets, particularly for MEGs where the overlap between datasets was strikingly larger (Fig. 3 and Supplemental Figure S4). In maize, consistent reanalysis identified a slightly increased number of imprinted genes shared between studies both for MEGs and PEGs.

It was previously proposed that the discrepancies might at least in part stem from different data analysis pipelines to call imprinted genes and, in fact, filtering with similar filtering conditions produced more overlap (Gehring et al., 2011; Pignatta and Gehring, 2012). However, even after consistent reanalysis, a high number of imprinted candidate genes are unique to a single dataset, and reanalysis is able to increase the overlap across datasets only by a limited degree. This suggests that biological variation (such as different environmental growth condition, different developmental seed stage, stochastic allelic expression differences), technical differences (e.g., library preparation/complexity, batch effects [Wang and Clark, 2014; Castel et al., 2015]), and differences in sequencing depth inherent to the datasets contribute to these discrepancies and cannot be corrected for *in silico*.

Particularly in maize, reanalysis did not increase the concordance between datasets. Even though the developmental seed stages did vary to some extent (Table 1; 10-14 DAP) we cannot fully explain this finding by biological variation only. All original analyses used the same statistical test and highly similar conditions for allelic bias filtering. Possibly, the published original analyses of maize datasets used filtering conditions close to the optimum, such that edgeR-based reanalysis could not further improve concordance. Furthermore, the co-identification of imprinted genes is additionally hampered by unequal read coverage: 48 of 129 imprinted genes proposed by Zhang and colleagues (2011) had too few reads to characterize imprinted expression in the data of Waters and colleagues (2011). The relative importance of biological *versus* technical biases on the large discrepancy between the datasets from maize is currently difficult to assess.

### A Statistical Approach that Uses Generalized Linear Models and Takes Into Account Replicate Information Largely Increases Sensitivity

Our statistical approach involves modeling of count data and is quite different from the approach chosen by the authors in the original publications. Our approach relies on generalized linear models and edgeR, which is a commonly used, well-established and robust method for differential expression analysis of RNA-seq data. By interpreting allelic counts of a cross as separate samples and imprinting analysis as a differential gene expression problem, we benefit from the power and flexibility of generalized linear models based on edgeR. The method is highly flexible, i.e. also batch effects can be modeled. The general methodology used here is also applicable to imprinting analysis in other cell types where a deviation from 1:1 is tested. A further advantage of our statistical approach is that it circumvents somewhat arbitrary minimum cut-offs of allelic imbalance. Most genes are only partially imprinted and we believe that the current knowledge does not support the exclusion of genes with moderate allelic imbalance yet reaching statistical significance. Our approach also allows (nearly) genome-wide ranking of genes according to their likelihood of being imprinted, allowing downstream applications, such as gene set enrichment analysis, with increased statistical power (e.g. for Gene Ontology analysis).

Prior to read counting, many steps are required to map, filter, and count allelic reads. To assess the importance of the preprocessing/counting relative to statistics, we reanalyzed the original allelic counts from Gehring et al. (2011) using our method. We found a large discrepancy for MEGs and PEGs with original gene lists, whereas MEGs and PEGs largely agreed with the gene lists obtained from complete reanalysis with our analysis pipeline (data not shown). Given the large discrepancy, the statistical approach seems to have a larger relative contribution than the earlier steps required to obtain the variant counts.

### Simulation Reveals that edgR-based Analysis Outperforms Other Methods

The edgeR-based approach performed well in our simulation setting and clearly outperformed the other tested methods. Notably, the other methods based on Fisher’s exact test seem overly conservative and missed most spike-in imprinted genes. edgeR was the only method to predict false positives due to a relatively poor FDR control for the lowest quarter of counts (10-23 counts per gene) where 4 of edgeR’s totally 8 false positive imprinted genes were identified. edgeR’s specificity could be further increased by selecting a larger minimal number of counts per gene (e.g., a cut-off of 20 allelic counts per gene).

Importantly, our simulation did not aim to model biological variability predicting biological replicates. All biological systems have inherent biological variation and edgeR can account for it, whereas Fisher’s exact test completely ignores the within-condition variability, as it requires counts from replicates to be summed up for each condition. Therefore, we expect that edgeR would outperform methods using Fisher’s exact tests even more strikingly when biological replicates are available.

### Higher Coverage, Replicate Samples, and edgeR-based Analysis Could Improve the Identification of Imprinted Genes

The first generation of genome-wide studies identified many new imprinted genes in plants, yet a considerable proportion of genes could not be characterized due to low sequencing depth or insufficient genetic heterogeneity between the parents. Future experiments in genomic imprinting will be performed using paired-end reads with high sequencing coverage. With the rapid development in NGS, higher coverage is now readily achievable, which will notably increase the statistical power to detect imprinted genes. In the studies reanalyzed here, low counts were observed for many genes close to the minimal coverage cut-off of 10, where the variance is large and a large allelic bias is required to reach significance. Importantly, a sufficient number of biological replicates (i.e., at least three per reciprocal cross [Liu et al., 2014]) would permit to reliably estimate the variability from the data, enabling the performance of a more robust differential expression analysis and a more reliable estimation of the total number of imprinted genes.

### Biological Perspectives

When comparing datasets from different studies, we assume that the same set of genes is imprinted over the sampling time period. Partial or complete violations of the assumption also decrease the amount of overlap across datasets in addition to technical and analytic biases. Indeed, first studies showed dynamic expression of imprinted genes in maize (Xin et al., 2013). Furthermore, we cannot solely rely on statistical significance in calling a gene imprinted or not. There are several ways to prioritize a gene list after statistically calling genes with an allelic bias in a given tissue. First, filtering the gene lists by fold difference between the two parental alleles could help to identify genes that can more easily be validated experimentally. Second, filtering the gene lists against tissue-specific expression data from seeds might identify genes relevant to the tissue of interest (Belmonte et al., 2013; Schon and Nodine, 2017). However, expression of a given gene in other (maternal sporophytic) tissues does not exclude an allelic bias in the fertilization products (Raissig et al., 2013). Third, comparing the gene list against central cell and sperm cell expression data (Borges et al., 2008; Wuest et al., 2010; Schmid et al., 2012) can inform to what extent the allelic bias is a result of expression in the fertilization products or might at least partially represent carry-over of gametic transcripts that were produced prior to fertilization.

An improved assessment including larger gene sets (by using different sets of strains), as well as imprinted genes with moderate allelic imbalance, will provide further insights into the extent and biological significance of genomic imprinting. Having the full catalog of imprinted genes in several plant species will also allow the tackling of evolutionary questions about genomic imprinting, including its origin and fixation, the conservation of imprinted genes, and their gain and loss of imprinting status on the phylogenetic tree.

Lastly, we have to stress that without further confirmation, gene lists are not representative with regard to the absolute number of imprinted candidate genes expressed in the endosperm. Considering this, it is indispensable to confirm bioinformatically identified imprinted candidate genes by alternative methods, such as allele-specific expression analysis using RT-PCR and Sanger sequencing, pyrosequencing, and/or reporter gene assays.

## MATERIALS AND METHODS

### Read Mapping and Counting

FASTQ-formatted raw reads were downloaded from the NCBI Short Read Archive (SRA) for endosperm experiments for *Arabidopsis* (GSM674847, GSM674848, GSM756822, GSM756824, GSM607727, GSM607728, GSM607732, GSM607735, GSM1276498, GSM1276500, GSM1276502, GSM1276504, GSM1276505, GSM1276508, GSM1276509, GSM1276512, GSM1276514, GSM1276515) and for maize (SRX105679, SRX105678, SRX114629, SRX114630, SRX047539, SRX047544, SRP031872, GSE48425). Hsieh samples obtained through laser capture microdissection were not included in the analysis as their inclusion reduced the overlap with other datasets (data not shown). Reads were quality-checked with the FastQC application (http://www.bioinformatics.babraham.ac.uk/projects/fastqc/). To reduce the bias in mapping reads towards reference alleles, we aligned the reads to a masked reference genome, in which bases at known polymorphic sites were replaced with “N”. Reads were mapped to the genome using STAR v2.3.0e (Dobin et al., 2012). Only reads that mapped to a unique position in the genome were considered for further analysis.

For *Arabidopsis* the genome annotation TAIR10 (http://www.arabidopsis.org/) was used, and for maize AGPv3 annotated genes were downloaded from Ensembl Plants Release 31 (http://plants.ensembl.org/ [Monaco et al., 2013]). *Arabidopsis* SNP annotation files were obtained from the 1001 Genomes project (Cao et al., 2011; http://1001genomes.org/data/MPI/MPISchneeberger2011/releases/2012_03_14/). Maize hapmap v3.2.1 SNP variants (Bukowski et al., 2015) in VCF format (Release date 3/3/2016) were obtained from the Panzea database (http://cbsusrv04.tc.cornell.edu/users/panzea/download.aspx?filegroupid=15) and filtered for high-confidence (flag LLD) biallelic SNP variants which were polymorphic between B73 and Mo17 and homozygous in both inbred strains. Allelic reads were counted at previously identified SNP positions between homozygous parentals, using python v3.2 with pysam v0.8.4 (Li, 2011), while positions with low sequencing quality (phred quality <20) were excluded. No more than one SNP was counted per read to prevent pseudo-replication. Counts were summed up per gene and variants with less than 10 reads (summed across the two reciprocal crosses) were discarded.

### Testing for Allele-specific Expression

To assess allele-specific expression, we used edgeR version 3.4.2 (Robinson et al., 2010). It uses an empirical Bayes estimation based on the negative binomial distribution. For library size normalization and to eliminate composition biases between libraries we used the TMM (Trimmed Mean of M-values) method. TMM normalization keeps the ratio between maternal and paternal allelic reads in a cross at approx. 2 (data not shown). Experiments were analyzed using a generalized linear model with a paired design (the two allelic counts of the same cross treated as paired samples) and at least two biological replicates (the two reciprocal crosses). When biological replicates were available, they were included in the model. We used tagwise dispersion estimates. False Discovery Rate (FDR) was calculated according to Benjamini and Hochberg (1995). The R software version 3.0.3 (R Core Team, 2012) was used for statistical analysis and for creating graphs.

### Simulation of genomic imnprinting

For generating synthetic data we used the function makeExampleDESeqDataSet of the DESeq2 version 1.2.10 (Anders and Huber, 2010) R/Bioconductor package. Mean and dispersion parameters that were used in the simulation were estimated from real RNA-seq data (interceptMean=2, interceptSD=3). Two biological replicates were simulated to serve as the two reciprocal crosses. No outlier counts and differential expression were introduced. The total number of genes in each simulated dataset was 15,000, and their true proportion of maternal reads was set to 2/3. Then we randomly picked 200 genes and modified their imprinting status, 50 each of strong MEGs (99% maternal reads), weak MEGs (85%), strong PEGs (34%) and weak PEGs (48%). Random allelic read sampling was modeled by sampling from a binomial distribution with the probability of success set to the true proportion of maternal reads.

We evaluated three methods for identifying genes with parent-of-origin specific expression: edgeR, Fisher’s exact test and Stouffer’s method. Fisher’s exact test does two separate statistical tests for the two reciprocal crosses and the resulting two p-values per gene were combined with Stouffer’s method (calculated by sumz function of R package metap). Benchmarking was performed using Venn diagrams and Receiver Operating Characteristic (ROC) curves with iCOBRA (Soneson and Robinson, 2016) https://github.com/markrobinsonuzh/iCOBRA.

### Comparison with Published Gene Lists

Published gene lists were compiled from the Supplementary Data of the respective publications. For the Waters B73xMo17 comparison, the updated list from Waters et al. (2013) was used, not the original one (Waters et al., 2011). If lists of imprinted genes were available at various stringencies, we used the least stringent list (e.g. moderately imprinted genes for the Waters dataset). Accession/inbred-specific imprinted genes were omitted. Only the lists of 106 MEGs and 91 PEGs at 12DAP (Prof. Xiaomei Lai, personal communication) described in Zhang et al. (2014) were used as the endosperm samples at 10DAP were already described in Zhang et al. (2011).

### Saturation Plots

In order to assess the degree of undersampling, we performed random subsampling on the count data and performed the same processing and statistical analysis steps as for the full data.

### Code and data availability

Scripts and data used in this manuscript are available on github (http://www.github.com/swyder/Reanalysis_plant_imprinting).

## ACKNOWLEDGEMENTS

We thank Malgorzata Nowicka and Prof. Mark Robinson (Institute for Molecular Life Sciences, University of Zürich), as well as Marc W. Schmid (Institute of Plant and Microbial Biology, University of Zürich) for helpful discussions. **This work was supported by the University Research Priority Program ‘Evolution and Action’ of the University of Zürich and, in part, by an Advanced Grant of the European Research Council (Nr. 243996) to U.G.**

## SUPPORTING INFORMATION

**Supplemental Table S1.** List of candidate imprinted genes identified in this study for *Arabidopsis*. Genes are sorted with decreasing probability of being imprinted in Gehring, Hsieh and Pignatta datasets.

**Supplemental Table S2.** List of candidate imprinted genes identified in this study for maize.

**Supplemental Table S3.** Jaccard similarity indices between originally published datasets or after reanalysis using edgeR or Stouffer’s method in *Arabidopsis* and maize. The same numbers of topmost imprinted genes were selected from the reanalyzed datasets.

## SUPPLEMENTAL DATA

**Supplemental Figure S1.**
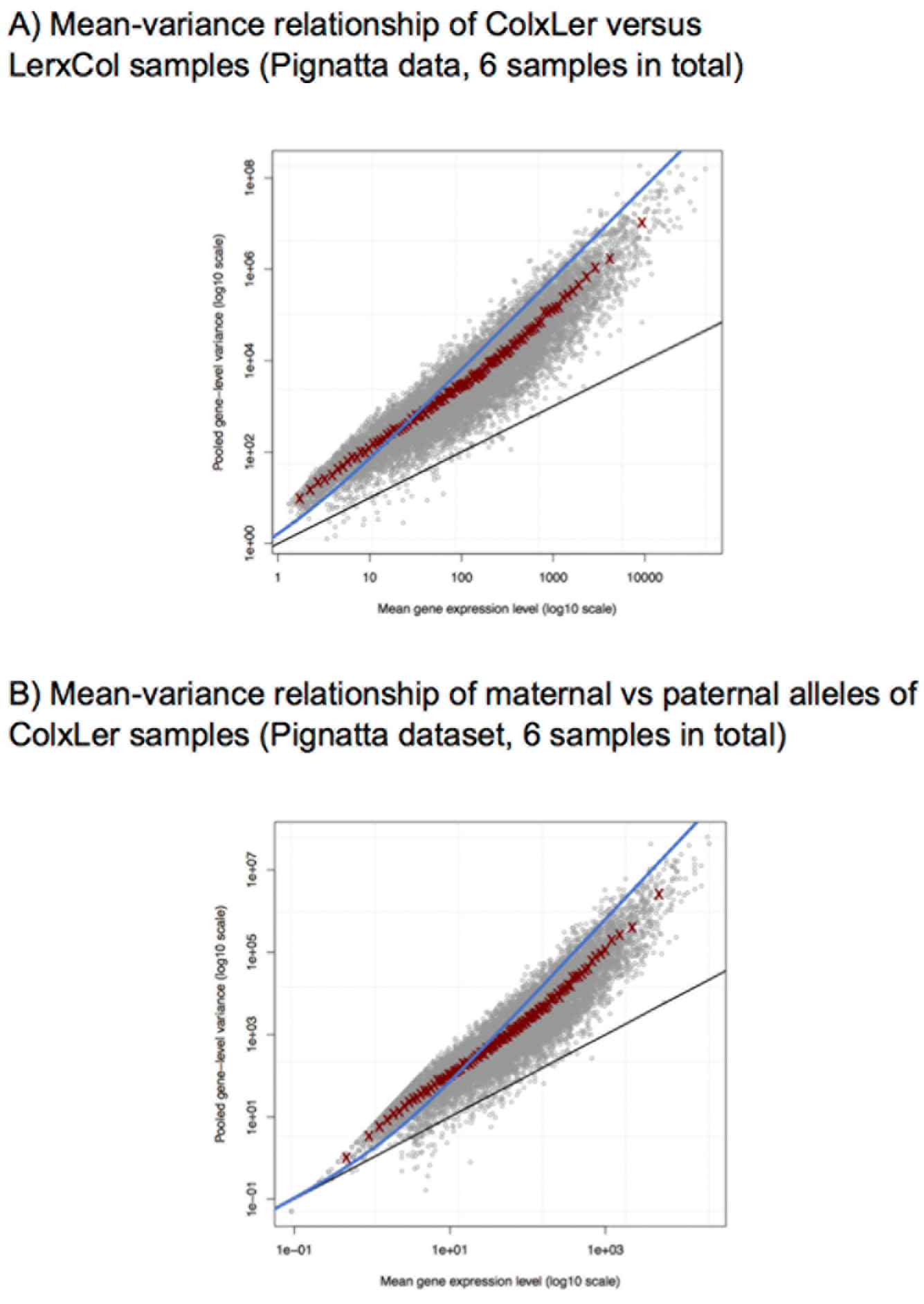
Count overdispersion in the Pignatta dataset. Mean and variance are plotted for A) ColxL*er* versus L*er*xCol samples (3 samples each) and B) maternal versus paternal samples in ColxL*er* crosses (3 samples each). The blue line shows the Negative Binomial model with common dispersion and the black line shows the Poisson mean-varieance relationship. The best fit with the data is observed by averaging raw variance for tags split into bins by overall expression level (red crosses). In both plots the variance for the counts between samples is much larger than the mean, indicating overdispersion.

**Supplemental Figure S2.**
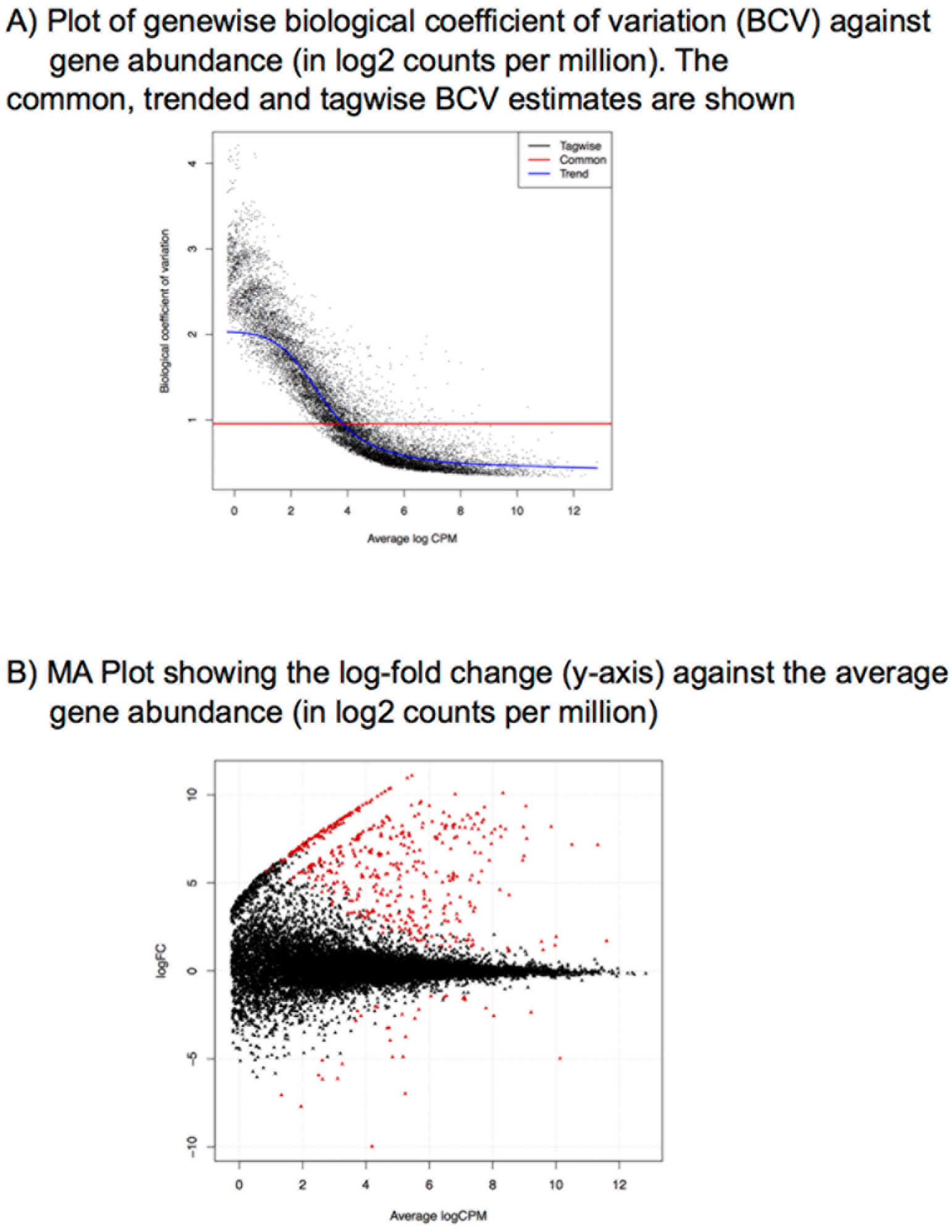
Diagnostic plots for the Pignatta dataset. A) Genewise biological coefficient of variation (BCV) against gene abundance (in log2 counts per million). B) MA plot of the Pignatta dataset. Genes with a significant allelic bias at a 5% FDR cut-off are highlighted in red.

**Supplemental Figure S3.**
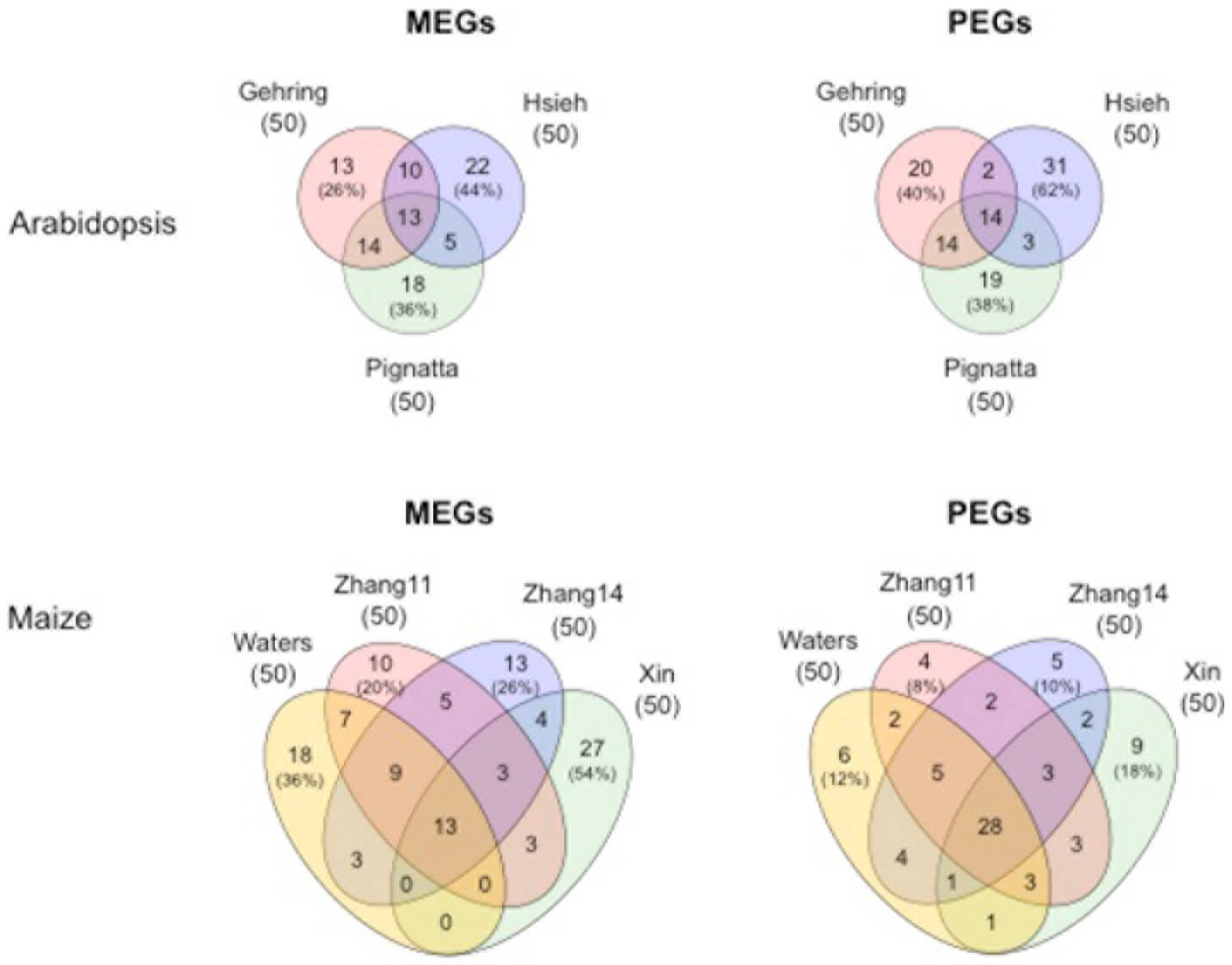
Venn diagrams showing the overlap between the top50 imprinted candidate genes across datasets when reanalyzing the raw data using the same standardized method, using generalized linear models and edgeR. Numbers in brackets denote the percentage of non-shared genes relative to the full set detected in the dataset.

**Supplemental Figure S4.**
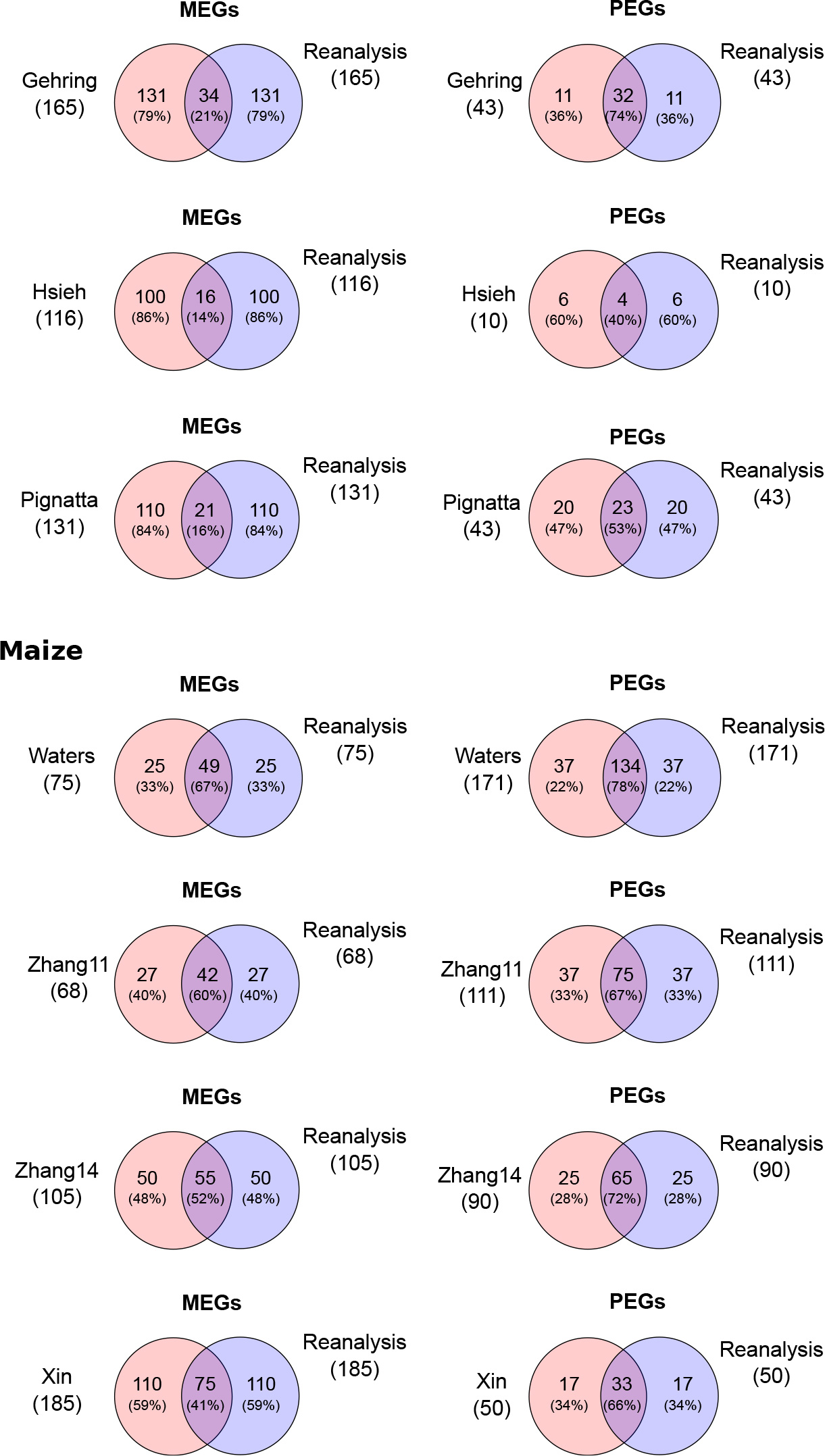
Pairwise comparison of imprinted genes between the originally published analysis and the reanalysis using generalized linear models and edgeR. The same numbers of topmost imprinted genes were selected from the datasets reanalyzed with generalized linear models and edgeR. Numbers in brackets denote the percentage relative to the full set.

**Supplemental Figure S5.**
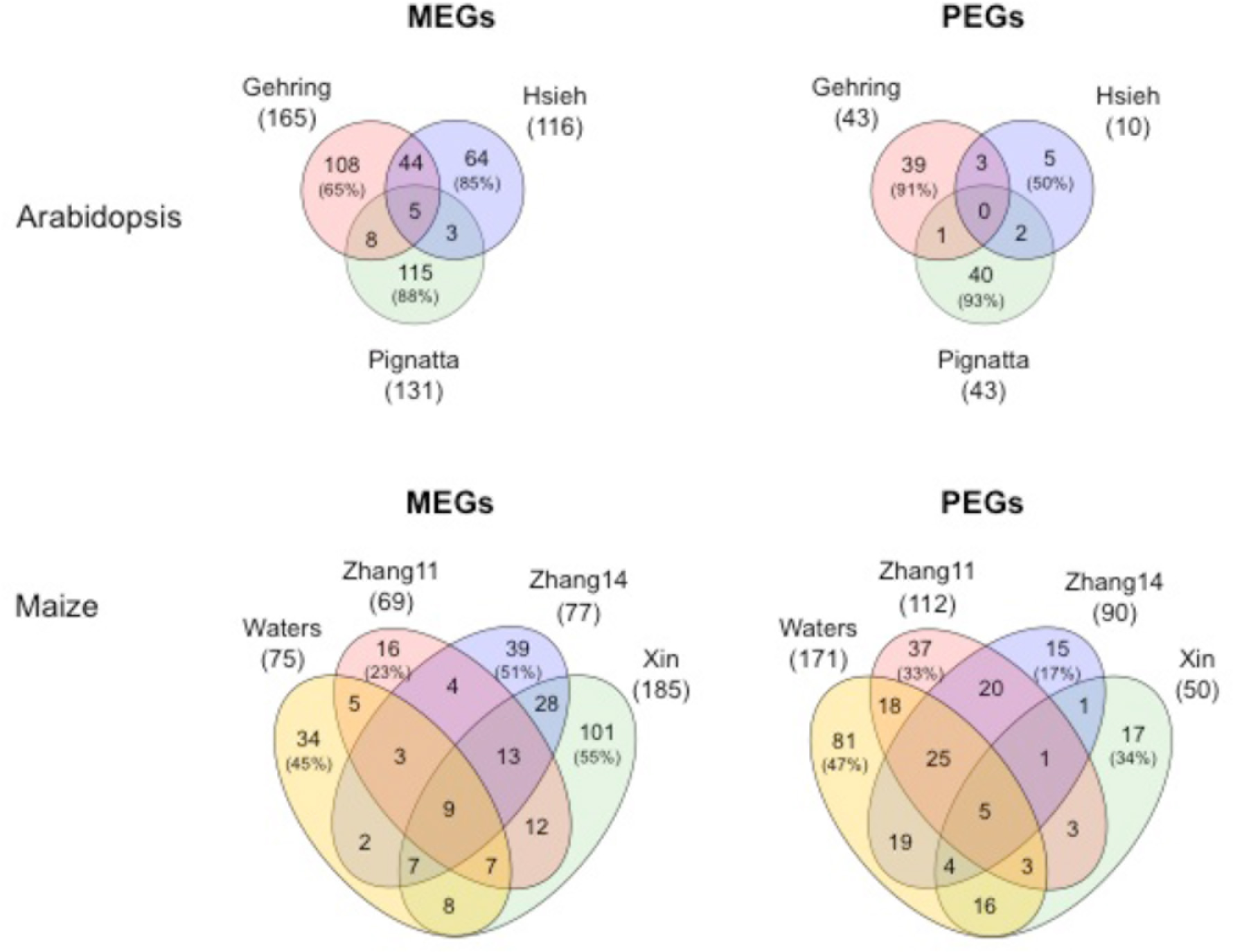
Venn diagrams showing the overlap between imprinted candidate genes across datasets when reanalyzing the raw data using the "Fisher-combined" (Stouffer’s) method. Numbers in brackets denote the percentage of non-shared genes relative to the full set detected in the dataset.

**Supplemental Figure S6.**
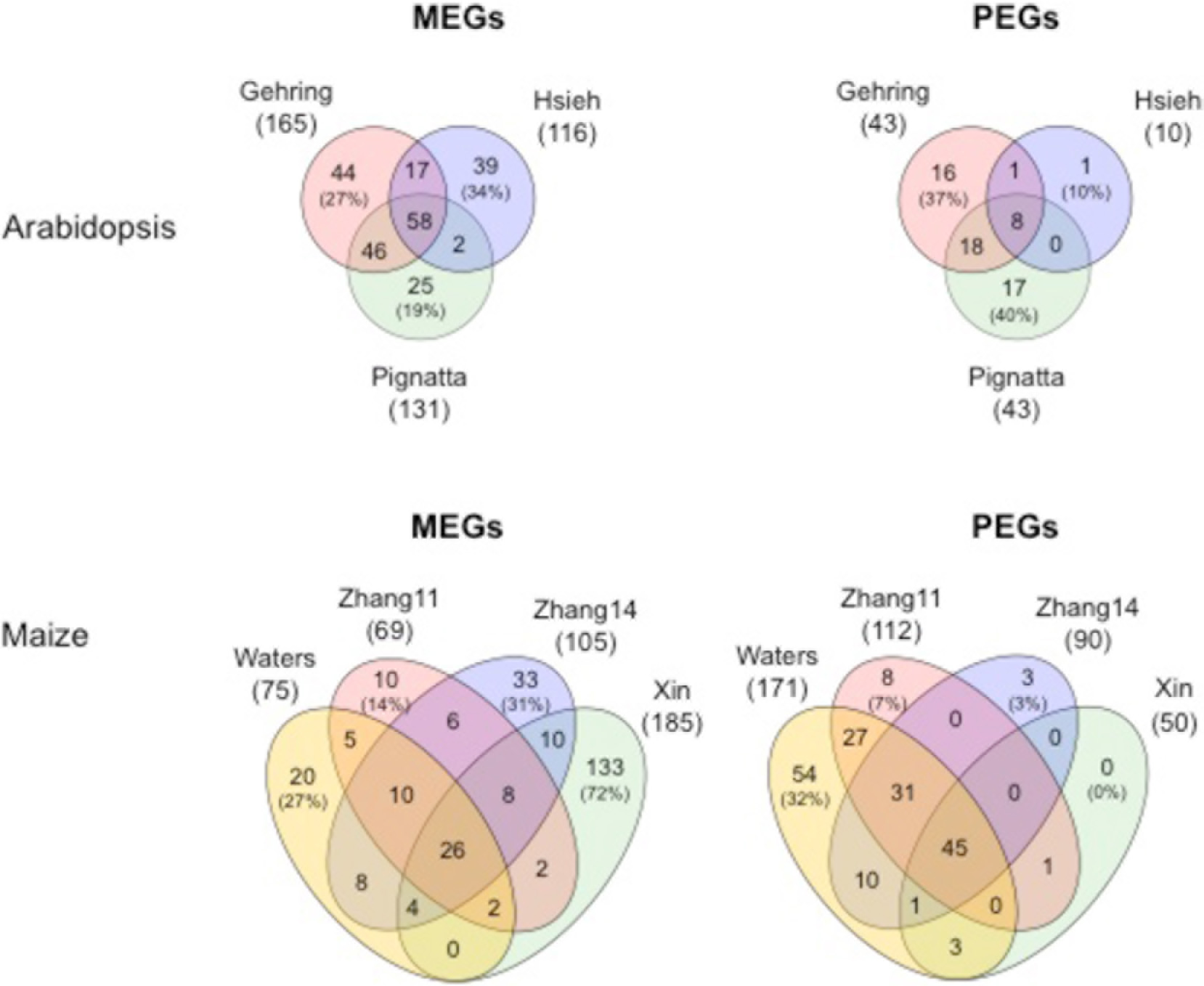
Venn diagrams showing the overlap between imprinted candidate genes across datasets when reanalyzing the raw data using generalized linear models and edgeR. Numbers in brackets denote the percentage of non-shared genes relative to the full set detected in the dataset.

